# Evaluating apoptotic gene efficiency for CHO culture performance using targeted integration

**DOI:** 10.1101/2024.05.29.596411

**Authors:** David Catalán-Tatjer, Saravana Kumar Ganesan, Iván Martínez-Monge, Lise M. Grav, Jesús Lavado-García, Lars K. Nielsen

**Author notes:** Corresponding author: Jesús Lavado-García.

## Abstract

Chinese hamster ovary (CHO) cells have long been the favoured platform for producing complex biopharmaceuticals such as monoclonal antibodies (mAbs). Cell death is a critical factor in all CHO cultures, dictating duration until harvest in batch cultures and viable cell density in perfusion. The programmed cell death, or apoptosis, pathway has been widely studied due to its relevance in affecting cell culture performance and the extensive knowledge about its protein-to-protein interaction network. However, clonal variation seen with random integration has confounded results and it remains unclear which effector genes should be overexpressed. Here, we employed the recombinase-mediated cassette exchange (RMCE) strategy to develop isogenic cell lines expressing one copy of erythropoietin, as model protein product, and various anti-apoptotic genes: bcl-2 from CHO and human origin, bcl-xL from CHO and human origin, mcl-1 and bhrf-1. We tested the generated isogenic cell lines in the presence of sodium butyrate, a well-known apoptotic initiator, in batch culture. The most promising candidates were cultured in fed-batch in the microbioreactor ambr®15 system. The observed phenotype varied significantly depending on the overexpressed gene, therefore the metabolic differences were further characterized using multiplexed quantitative proteomics. We showed that overexpressing bcl-2 from CHO origin significantly improved productivity and established a methodology to successfully test candidate genes via targeted integration. This will enable future metabolic engineering strategies to be more comparable and overcome the challenges faced thus far.

## INTRODUCTION

Programmed cell death or apoptosis represents the culmination of a signalling cascade activated during periods of heightened cellular stress, stemming from factors such as mechanical or oxidative stress, as well as nutrient depletion (Krampe & Al-Rubeai, 2010; Mohamed et al., 1998; R. Singh et al., 2019). The loss of cell viability ultimately dictates when bacth cultures are terminated, hence many studies have been dedicated to studying and delaying apoptosis as much as possible, following the guiding principle of “the longer the culture, the higher the productivity” (Rahimi et al., 2023; R. P. Singh et al., 1994). The first exponent of this idea was the introduction of the fed-batch culture, in which limiting nutrients were supplemented while the culture was already growing and producing, aiming to delay the apoptosis triggered by the absence of these key nutrients (Bibila & Robinson, 1995; Ochoa, 2016). Perfusion has been used to extend cultures indefinitely (Bielser et al., 2018) and achieving record-high cell density, constrained only by nutrient and oxygen supply to the culture (Clincke et al., 2013; O’Flaherty et al., 2020). Even perfusion, however, is affected by apoptosis with steady state cell density at low bleed rates dictated by the death rate.

Apart from feeding strategies to mitigate nutrient depletion (Altamirano et al., 2000), numerous scientists have explored various genetic engineering approaches to delay apoptosis. Recently, CHO cells were engineered to overexpress or knock out apoptotic-related genes (Grav et al., 2015; MacDonald et al., 2022). As reviewed by Henry *et al*. Bcl-2 family proteins, which include Bax, Bak, Bcl-2, Bcl-xL and Mcl-1 have been the most studied regarding apoptosis delay (Henry et al., 2020). Bax and Bak function as effector proteins capable of inducing pores on the mitochondrial membrane, thereby initiating apoptosis. Conversely, Bcl-2, Bcl-xL and Mcl-1 serve as inhibitor proteins, impeding the generation of these pores (Henry et al., 2020). Many publications report that overexpressing either Bcl-2, Bcl-xL, Mcl-1 or a combination of them, led to higher final titers as apoptosis was significantly delayed (Fussenegger et al., 2000; Majors et al., 2009). Other studies report on knocking out Bax and/or Bak to prevent apoptosis with promising results (Grav et al., 2015; Xiong et al., 2019). Furthermore, considering the impact of these proteins on the mitochondrial membrane and, consequently, their influence on the efficiency of the electron transport chain and the tricarboxylic acid (TCA) cycle, some researchers have evaluated these cell lines with regard to these specific pathways with interesting results (Templeton et al., 2014).

Regarding the success of these strategies and the development of advanced techniques for cell line generation, scientists have tried to replicate and improve previously reported data, with contradicting results as compiled by Henry *et al*. (Henry et al., 2020).

For instance, while some authors describe an increase in the specific productivity (q_p_) when overexpressing bcl-xL (Majors et al., 2008), other studies conclude that there was no significant change (Han et al., 2011). Similar discrepancies have been reported with bcl-2. Additionally, not all publications regarding the improvement in the duration of the culture agreed on the beneficial effects on the expression of either bcl-2 or bcl-xL.

This lack of reproducibility and discrepancies in the reported works might be attributed to clonal variation. The use of random integration to generate stable cell lines (Fussenegger et al., 2000; Ko et al., 2018; Mastrangelo et al., 2000) is associated with large clonal variation and it is uncommon to characeterize tens of strains to accommodate for this variation. Conversely, the study of cell pools suffer from a diffuse signal, particularly if exposed to extended batch culture. This concern was already highlighted by Henry et al. who concluded that establishing isogenic cell lines in a stable genomic location may be the only viable option to ensure meaningful comparisons (Henry et al., 2020).

In this study, we used our isogenic cell line strategy (Grav et al., 2018) to evaluate the anti-apoptotic genes bcl-2 and bcl-xL from human origin, bcl-2 and bcl-xL from CHO origin, mcl-1 and the apoptosis regulator Bhrf-1 from Epstein-Barr virus. Targeted integration was used to develop isogenic cell lines expressing one copy of each relevant gene to overcome clonal variation. In order to test the hypothesis related to productivity, one copy of erythropoietin (EPO) was integrated using the same strategy.

## RESULTS AND DISCUSSION

### Successful generation of stable isogenic subclones co-expressing EPO and anti-apoptotic genes

Clones generated via recombinase-mediated cassette exchange (RMCE) share the same genome and can therefore be termed isogenic. Consequently, we do not anticipate any significant differences between isogenic clones when assessing cell growth, metabolism, and production (Lee et al., 2015). This enables meaningful comparisons regarding the effects of gene expression, as it ensures that each gene is present in one copy under the same promoter and in the same gene location across all cell lines.

A CHO master cell line with an already integrated single landing pad previously generated by our group (Grav et al., 2018) was used to create the different isogenic cell lines harbouring anti-apoptotic genes (Figure 1).

**Figure 1.**
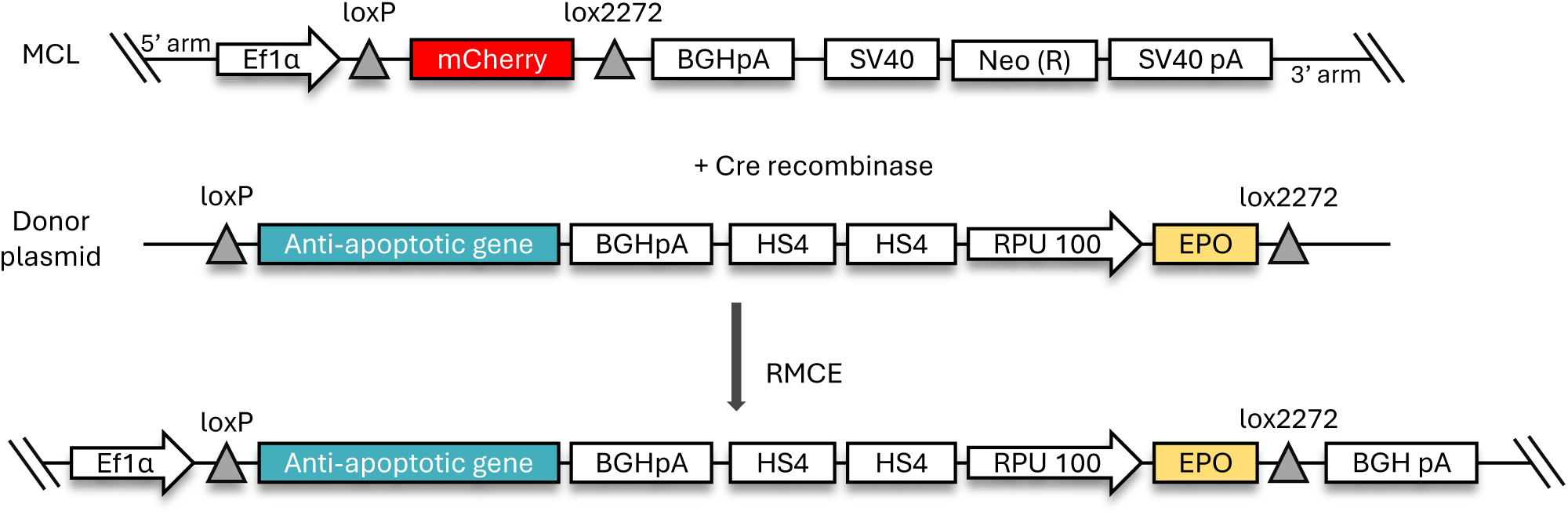
Isogenic cell line generation strategy. CHO-S master cell line (MCL) was transfected with two plasmids: a Cre recombinase plasmid and a donor plasmid. The donor plasmid was designed with the corresponding anti-apoptotic gene followed by the bovine growth hormone polyadenylation signal (BGHpA) and by two copies of chicken β-globin locus control region hypersensitive site 4 (HS4) which act as an insulator to enhance erythropoietin (EPO) expression. RPU 100 was selected as a promoter for EPO. LoxP sequence was added on the 5’ arm of the construct whereas lox2272 was located at 3’ arm. After dual transfection, cells were sorted based on mCherry fluorescence. Successful recombinants lost mCherry expression and where single cell sorted in 384-well plates.

We successfully generated and verified a minimum of three clones for each cell line co-expressing one copy of the anti-apoptotic gene and one copy of EPO. All donor plasmids employed in this step are described in Supplementary Figure S1.

The process of clone validation began with a junction PCR to verify the expected size of the inserted construct. Recognizing the potential for recombination to occur at other genomic sites, a subsequent step was required to detect any additional recombination events. Therefore, qPCR was utilized to assess the number of EPO copies present in the entire genome. Clones that underwent positive junction PCR validation and possessed a single copy of EPO (Supplementary Figure S2) were banked and characterized. A negative control was generated with the same methods replacing the anti-apoptotic gene with 3 STOP codons in the donor plasmids (Supplementary Figure S1).

A positive control was generated by knocking out the bcl-2 family effector protein, Bax and Bak, which are pivotal in pore formation in the mitochondrial membrane triggering apoptosis. The knockouts were performed in the negative control cell line and successfully confirmed by Western blot (Supplementary Figure S3).

### Anti-apoptotic isogenic cell lines were able to withstand the toxic effect of sodium butyrate

We studied the performance of the developed cell lines with sodium butyrate (NaBu) as it is commonly used in cell cultures to boost productivity, but will induce apoptosis at elevated concentrations (Joon et al., 2008; Wang et al., 2002). Therefore, a cell line able to better withstand the negative impact of NaBu while keeping cells viable for a longer time will be industrially relevant. For that reason, we examined the culture progression of three isogenic clones each for CHO-S mcl-1, bcl-xL CHO, bcl-2 CHO, bcl-2, and positive control BaxBak KO compared to the negative control upon incubation with 20 mM NaBu (Figure 2A). All bcl-2 genes except bcl-2 CHO significantly delayed cell death compared to the negative control (Figure 2B) with comparable performances on the viable cell density (VCD) progression (Figure 2C). As expected, the positive control BaxBak KO showed the greatest degree of protection.

**Figure 2.**
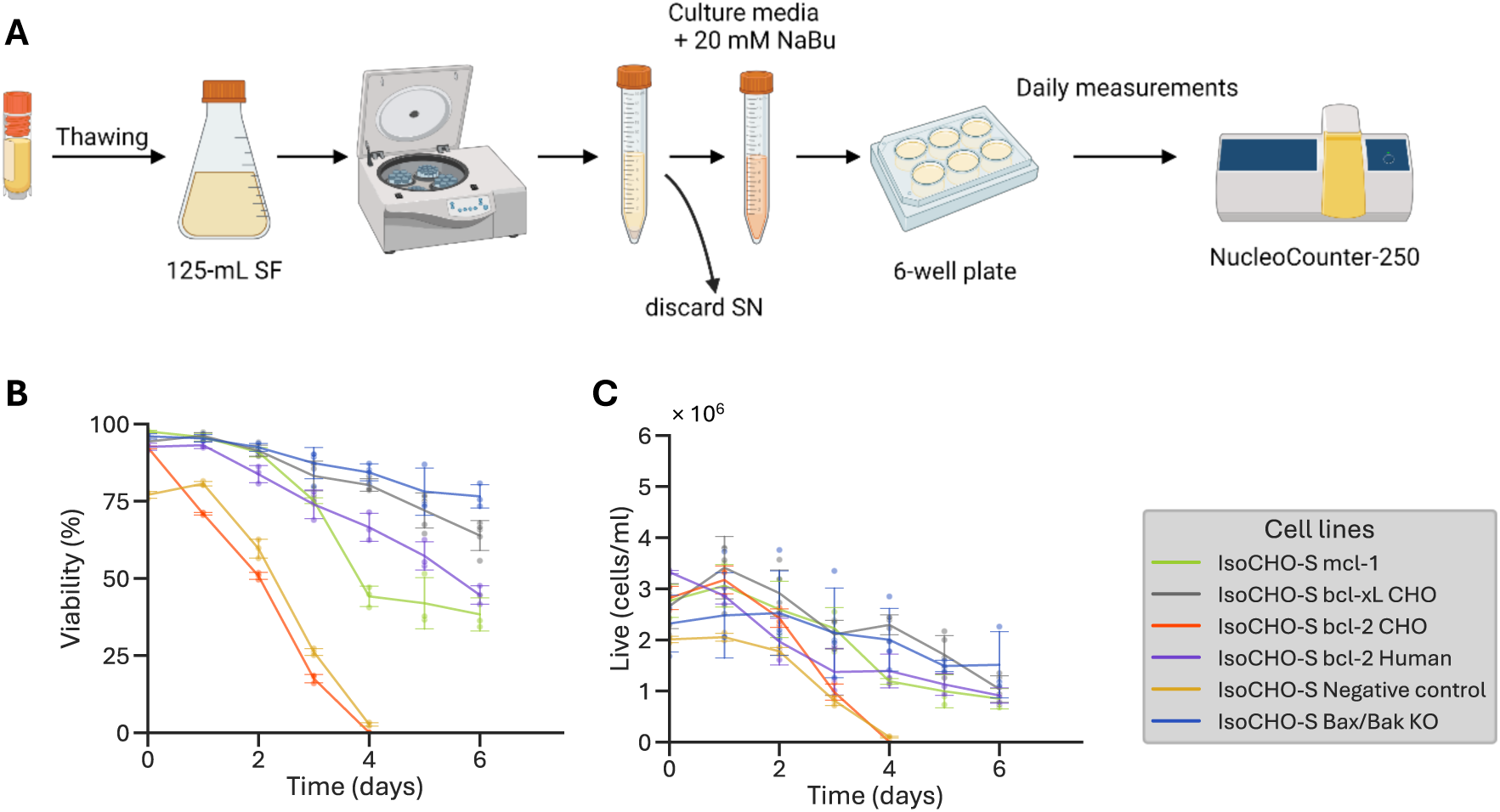
A) Cells were carefully thawed transferring the content from a cryovial containing 10 × 10^6^ cells to a 125-mL shake flask with a working volume of 30 mL. Then, cells were resuspended in culture mediasupplemented with 20 mM of Sodium Butyrate to reach a final concentration of 2.5 × 10^6^ cells/mL in 3 mL of working volume in a 6-well plate. (B) viability assessment and (C) cell counts were performed with the NucleoCounter-250.

The inability of native bcl-2 to suppress apoptosis was unexpected (Figure 2B). We first compared the cloned sequence with the native genes in Chinese hamster and human, to rule out a deleterious mutation in the CHO-S gene. The cloned gene was identical to the Chinese hamster gene (Uniprot ID Q9JJV8), which is homologous to the human bcl-2 alpha isoform (P10415-1), except for a three amino acid insertion in the latter. Of 25 single amino acid differences between Chinese hamster and human, 17 are identical with mouse instead (P10417). In proteomics (see later) bcl-2 CHO and human bcl-2 were quantified using identical peptides and their expression was comparable. We conclude that the relatively small changes in sequence could be responsible for the different profile observed.

This significant effect caused by a small genetic difference may explain inconsistent results in previous studies. Five out of thirteen previous experiments reviewed by Henry et al. observed no effect when bcl-2 was overexpressed while the remaining eight experiments reported an improvement of at least one day (Henry et al., 2020). This highlights the value of using isogenic cell lines when comparing genes capable of displaying subtle effects.

### Isogenic CHO-S bcl-2 CHO achieved the highest titre whereas CHO-S bcl-2 showed the longest batch culture time

While convenient, NaBu stress is artificial and does not capture the cellular stress observed in most cultures. We next compared the effect of overexpressing bcl-2 genes in conventional batch cultures over a span of 9 days. Two additional genes were included in the comparison: bhrf-1 is a homologous of bcl-2 originating from Epstein-Barr virus, while bcl-xL is the human version of bcl-xL allowing another comparison between human and CHO bcl-2 family genes.

Apart from CHO-S bhrf-1, all cell lines – including positive and negative controls – displayed similar development in batch until day 6 (Figure 3). While the maximum cell density achieved varied somewhat between cell lines (Figure 3A), this was countered by small differences in cell diameter (Figure 3C) and the metabolic profiles were near identical with depletion of glutamine at day 4 (Figure 3D) and glucose at day 5 (Figure 3E). From day 6 to day 7, VCD and viability of bcl-2 CHO experienced a precipitous decline, reducing viability by 50%, distinguishing it as the poorest-performing cell line (Figure 3B), mirroring its performance upon NaBu addition. Conversely, the human bcl-2 showed the slowest decline. CHO-S bhrf-1 achieved the highest cell density (Figure 3A) and produced the lowest amount of by-products lactate (Figure 3D) and ammonia (Figure 3E). However, the decline in viability was almost as fast as CHO-S bcl-2 CHO (Figure 3B).

**Figure 3.**
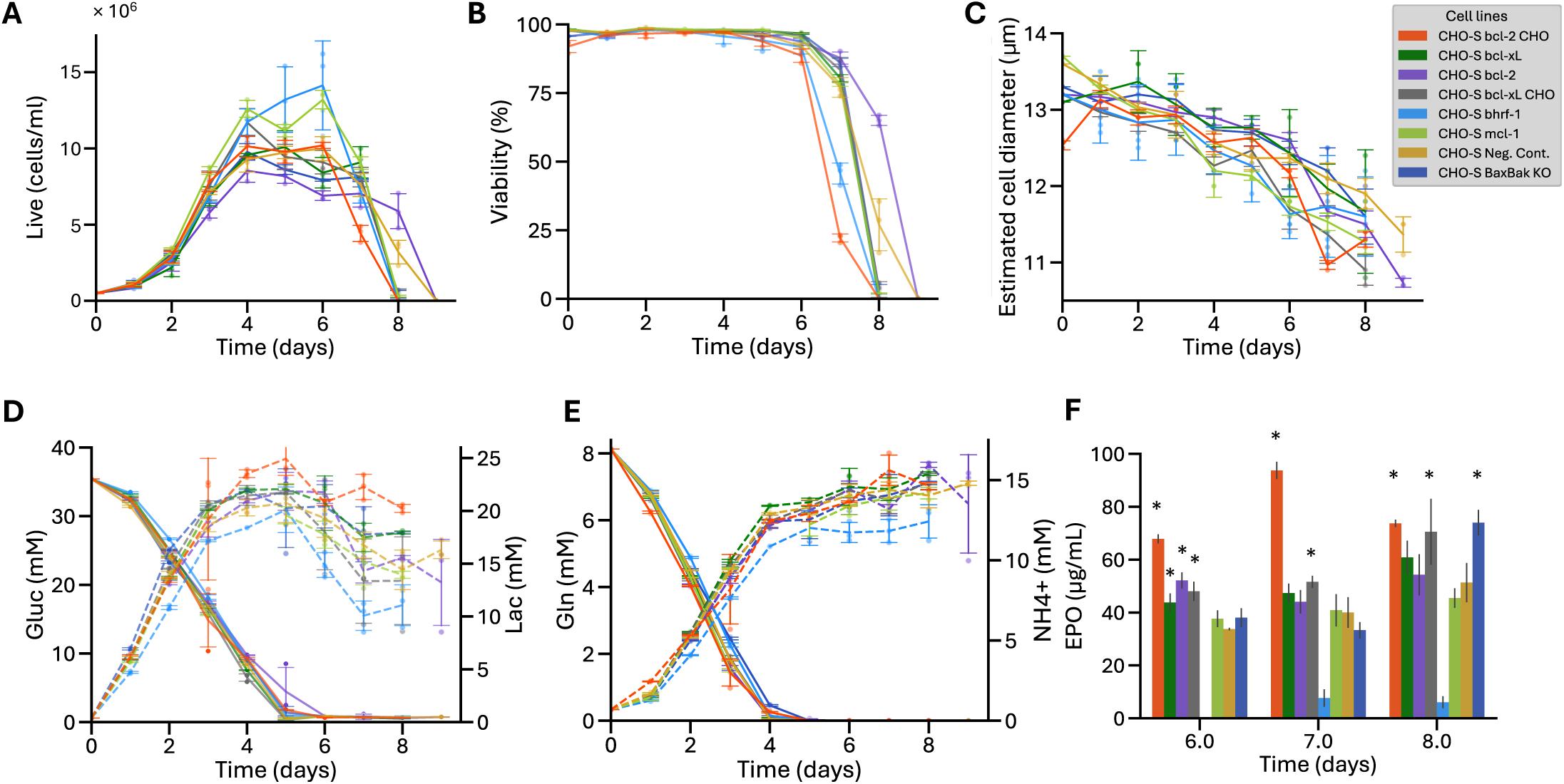
Batch culture of the studied isogenic cell lines. Development overtime of: (A) viable cell density in cells/mL, (B) viability in %, (C) estimated cell diameter in µm, (D) glucose and lactate in mM and (E) glutamine and ammonia in mM. (F) shows EPO concentration at day 6, 7 and 8. Statistically significant values compared to the negative control with a Dunnett’s test are (p-value<0.05) illustrated with “*”.

In terms of EPO production, both human and CHO bcl-2 and bcl-xL expressing cells achieved higher titre at Day 6, where mcl-1 and the two controls showed a lower titre (Figure 3F). On Day 7 and 8, the two CHO genes bcl-2 and bcl-xL were significantly higher than controls, while the human equivalents were not. The BaxBak clone caught up by Day 8, while the bhrf-1 cell line showed almost no production at any time point. The three CHO-S bcl-2 CHO cell lines showed a big peak at Day 7, possibly as a result of release from dying cells.

While the use of isogenic cell lines facilitated the direct comparison of growth and metabolic effects of anti-apoptotic genes, productivity measures are more difficult to interpret given that titres are measured after glutamine and glucose were depleted.

### Fed-batch cultures presented different phenotypes

Fed-batch cultures are a widely adopted manufacturing strategy wherein nutrients are supplied to compensate for potential nutrient depletion resulting from cell growth. Consequently, prolonged processes may lead to the overaccumulation of toxic by-products such as ammonia or lactate. Drawing from the observations in the batch culture, where two interesting profiles emerged among the outliers - bcl-2 CHO exhibiting the highest production and bhrf-1 showcasing the highest VCD alongside unexpected metabolic profiles - we perform a fed-batch in the ambr®15 system with three bioreactors for each cell line under pH and temperature-controlled conditions (Figure 3A, Figure 3D, and Figure 3E).

During the exponential phase of cell growth (from inoculation to day 5), minimal differences were observed in terms of VCD and viability (Figure 4A, 4B). Hereafter, performances began to diverge, with bhrf-1 achieving the highest VCD while producing the lowest amount of lactate (Figure 4D). This agreed with the previously observed results in batch mode. Surprisingly, in terms of viability, no significant anti-apoptotic behaviours were observed, as the cell line with the highest VCD experienced the most rapid decline in viability due to, presumably, nutrient depletion. This was already anticipated by a transcriptional study in 2006 which showed that the expression of bcl-2 does not change during a fed-batch (Wong et al., 2006). Moreover, bhrf-1 produced lower lactate than the negative control. However, it has to be noted that high variability in the lactate development for bhrf-1 was observed due to some datapoints falling below the limit of detection.

**Figure 4.**
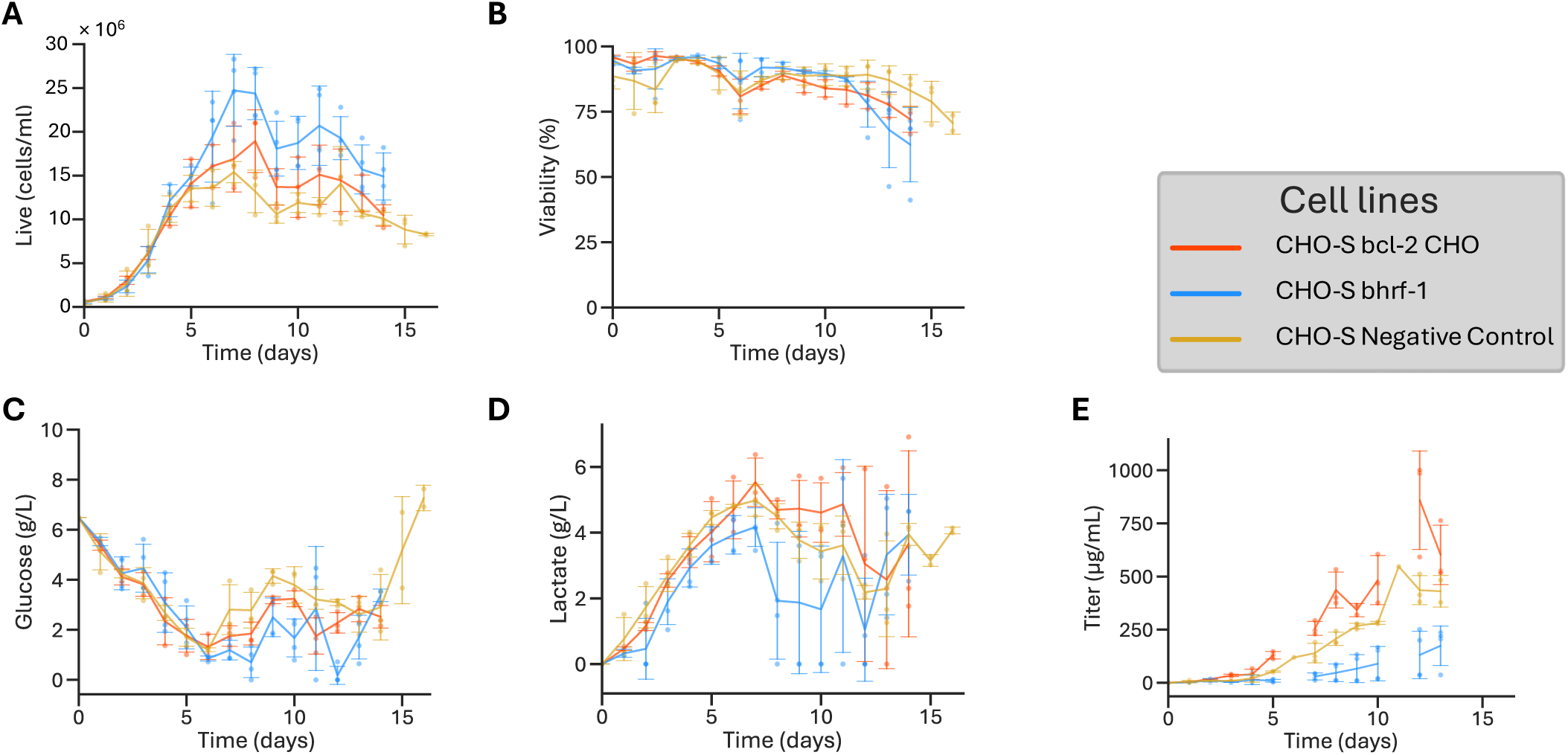
ambr15 fed-batch run for all studied isogenic cell lines. CHO-S bcl-2 CHO and CHO-S bhrf-1 alongside the CHO-S Negative Control cell line to study possible phenotype changes over the course of the process. (A) viable cell density in cells/mL, (B) viability in %, (C) glucose in g/L, (D) lactate in g/L and (E) EPO titer in µg/mL.

Interestingly, once again, bcl-2 CHO outperformed the negative control in terms of titer measured throughout the culture (Figure 4E), with similar glucose consumption (Figure 4C) and lactate production (Figure 4D) to the negative control.

Contrary to previously published data, which relied on shake-flask methods to conduct fed-batch experiments, we have, to the best of our knowledge, pioneered the assessment of bcl-2 CHO performance using the ambr®15 system. We chose this experimental setup due to its robustness, capability to operate reactors in triplicate, and its established track record in scalability. Throughout this project, our aim was to establish foundational insights into genetic comparisons, with a particular focus on anti-apoptotic cell lines. We believe that transitioning to equipment prioritizing reliability, provided it accommodates the experimental setup, is the optimal approach to achieve our goals.

In conclusion, the beneficial consequences of overexpressing anti-apoptotic genes were high VCD and increased titer for bcl-2 CHO.

### Relevant metabolic differences were observed between isogenic cell lines CHO-S bcl-2, bcl-2 CHO and bhrf-1 using proteomics

The different metabolic responses between cell lines warranted a closer investigation of the antiapoptotic responses. For this purpose, we used multiplexed quantitative proteomic analysis to compare bcl-2, bcl-2 CHO and bhrf-1 using the negative control as a reference. The apoptotic response was observed between late exponential phase (Day 3) and post depletion of glucose and glutamine but prior to major loss of cell death viability (Day 6) in batch culture (Figure 3; specific data points available in Supplementary Figure S4).

Only proteins identified with more than one peptide in the negative control on days 3 and 6 were selected for further analysis. We further restricted the analysis to proteins that displayed minimal change from Day 3 to 6 in the negative control (|ΔZq| < 1) to focus on proteins that were differentially changing in our studied cell lines and not in the negative control. Among the 3729 identified proteins, 2113 passed the cutoff of |ΔZq| < 1 (Figure 5A).

**Figure 5.**
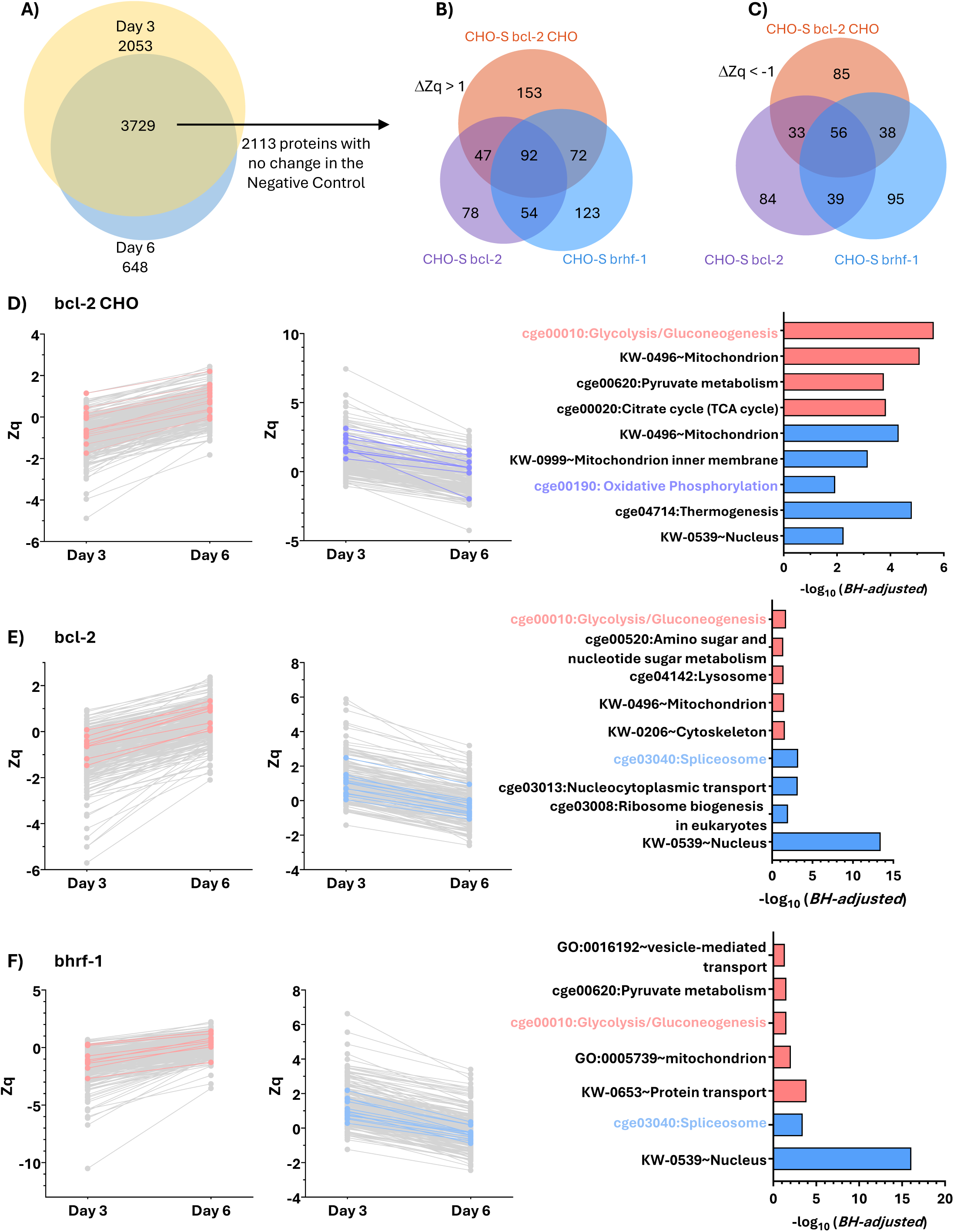
Metabolic differences in isogenic CHO-S bcl-2, CHO-S bcl-2 CHO and CHO-S bhrf-1. (A) Venn diagram of proteins identified with more than one peptide in the Negative Control at day 3 and at day 6. All proteins with a development of less than 1 unit of standardised variable, Zq, between day 3 and day 6 were excluded for further analysis. (B) Proteins with a ΔZq >1 from day 3 to 6 and (C) proteins with a ΔZq <-1 from day 3 to 6. (D) Shows all proteins from CHO-S bcl-2 CHO with (right) ΔZq <-1 from day 3 to 6 with proteins belonging to cge00190: Oxidative phosphorylation coloured in purple, (middle) ΔZq >1 from day 3 to 6 for cge00010:Glycolysis/Gluconeogenesis coloured in red and (left) relevant pathways selected based on logarithm of adjusted Benjamini score. Red bars indicate enrichment of upregulated processes and blue bars of downregulated process. (E) and (F) illustrate proteins filtered with the same criteria as Figure (D) for bcl-2 and bhrf-1 respectively with cge03040:Spliceosome proteins marked in blue.

From these 2113 proteins, 619 displayed an increasing trend (ΔZq > 1), while 430 demonstrated a decreasing trend (ΔZq < 1), in at least one of the modified cell lines. Notably, when visualizing the increasing and decreasing proteins in a Venn diagram, three distinct proteomes emerged (Figure 5B, Figure 5C), indicating that a significant number of proteins undergoing a specific trend were not shared among all cell lines. For instance, only 92 out of 619 showed the same increasing trend in all three cell lines, while 56 out of 430 proteins shared a decreasing trend. Consequently, we analysed each cell line individually using DAVID as an enrichment tool to identify processes that were enhanced in each cell line. Specific consumption and producing rates were calculated for the range of 3 to 6 days to further discuss some of the obtained enrichment results (Supplementary Figure S4).

In the case of bcl-2 CHO, increasing processes include glycolysis/gluconeogenesis (cge0010), mitochondrion (KW-0496), pyruvate metabolism (cge00620), and citrate cycle (cge00020) (Figure 5D). Out of the 13 overexpressed proteins belonging to the Kegg pathway of glycolysis/gluconeogenesis (cge0010), 3 participate in the citrate cycle and the pyruvate metabolism simultaneously (pyruvate dehydrogenase E1 component subunit beta (EC 1.2.4.1), acetyltransferase component of pyruvate dehydrogenase (PDH) complex (EC 2.3.1.12) and phosphoenolpyruvate carboxykinase (PEPCK) (EC 4.1.1.32)) (Supplementary Table S1). PDH upregulation suggests a lack of available ATP. PEPCK upregulation might be responsible for providing precursors for nucleotide and amino acid biosynthesis (Yu et al., 2021). This phenotype might explain some of the features observed in the metabolism of the bcl-2 CHO. It has been widely reported that when the glucose is depleted from the medium, cells start consuming lactate and, consequently, their metabolism shift towards pyruvate production (Kumar et al., 2021; Martínez-Monge et al., 2019; Martínez et al., 2013). This could be hinted by the overexpression of these proteins showing a flux coming out of the mitochondria to the cytosol which can benefit the biosynthesis pathway of different molecules such as amino acids (Yu et al., 2021), a pathway that has been positively enriched on bcl-2 CHO (Figure 5D). Pyruvate can still enter the TCA cycle and be converted to oxalacetate by pyruvate carboxylase as the proteomics analysis was based on relative quantification and the lack of overexpression indicates that the value is equivalent to the negative control, not that it is absent from the cell. The carbon flux moving towards gluconeogenesis and branching into these diverse biosynthesis pathways might explain the decrease rate of glucose consumption observed in bcl-2 (Supplementary Figure S4E).

Conversely, the oxidative phosphorylation (KW-0249), thermogenesis (cge04714), mitochondrion inner membrane (KW-0999), and nucleus (KW-0539) significantly decreased over time while they remained constant in the negative control (Adjusted BH < 0.05) (Supplementary Table S2). Considering these results, we hypothesized that the decrease in the oxidative phosphorylation pathway could reduce the availability of H^+^ inside the mitochondria, affecting the citrate cycle proton requirements enhancing the gluconeogenesis instead of the glycolysis. Considering how intertwined all the impacted pathways are and how a decrease of a flux can be compensated in an increase somewhere else, it is important to verify these hypotheses with the aid of more complex studies by, for instance, adding non-radioactive isotope tracers (usually synthetic metabolites labelled with carbon-13 (^13^C) (Coulet et al., 2022).

Proteins changing in bcl-2 cell line (Figure 5E) exhibited similar development to bcl-2 CHO in terms of glycolysis/gluconeogenesis (cge0010) being upregulated. The glucose consumption rate between day 3 and day 6 was indeed, statistically significantly larger than the negative control, showing a higher consumption of glucose and a higher lactate production (Supplementary Figure S4E and Supplementary Figure S4G). However, pyruvate metabolism (cge00620), and citrate cycle (cge00020) were not statistically significantly upregulated (Adjusted BH > 0.05). Interestingly, cytoskeleton (KW-0206) proteins were upregulated while nuclear (KW-0539) proteins were downregulated. This might explain why cell size in bcl-2 CHO is larger on day 6 compared to the negative control (Figure 3C). The decrease in spliceosome (cge03040) and ribosome biogenesis-related (cge03008) proteins is a phenotype related to the smaller growth observed in this cell line compared to the other one, as presented in Figure 3A. All results from enrichment analysis are grouped in Supplementary Table S3 and S4.

Lastly, proteins changing in bhrf-1 cell line showed upregulation in glycolysis/gluconeogenesis (cge0010) with 8 proteins and in pyruvate metabolism (cge00620) with 7 proteins (Figure 5F). The decrease of nuclear (KW-0539) proteins mostly translated to the decrease of spliceosome (cge03040) proteins. All up and downregulated processes are available in Supplementary able S5 and S6. The spliceosome is responsible for the removal of introns from nuclear pre-mRNA. However, considering the crucial role of the spliceosome in the proper function of the cell, many proteins are redundant (Wahl et al., 2009).

Interestingly, although the three modified cell lines all display an upregulation in the glycolysis/gluconeogenesis pathway, only 4 proteins were common in the upregulation: Phosphoglucomutase-2, S-(hydroxymethyl) glutathione dehydrogenase, multiple inositol polyphosphate phosphatase 1 and aldo-keto reductase family 1 member A1. These proteins are scattered in the pathway with no direct connection between them showing that, although the glycolysis/gluconeogenesis was significantly enriched, each cell line acted differently.

Another interesting detail is that most of the shared overexpressed proteins were localised in the cytoplasm (17 in cytosol, 13 in mitochondrion, 30 in cytoplasm unspecified) whereas most of the downregulated proteins were localised in the nucleus (15 proteins).

In conclusion, the overexpression of a single gene with apparently the same function but different origin (human, CHO, and viral) resulted in three distinctly different phenotypes. A proteomic analysis facilitated drawing conclusions to explain the different metabolic profiles.

## MATERIALS AND METHODS

### Cell culture

CHO-S cell lines (Thermo Fisher Scientific, Waltham, MA USA) were cultivated in disposable 125-mL polycarbonate shake flasks with a working volume of 30 mL. CD CHO medium (Gibco™, Buffalo, NY, United States) was supplemented with 8 mM of glutamine (Gibco™), 1 % of Anti-Clumping Agent (Gibco™) and 0.01 % of Antibiotic-Antimycotic 100X (Gibco™). Cell cultures were incubated in a 5 % CO_2_ air mixture at 37 °C in a Heracell 150 incubator (Thermo Fisher) and agitation of 120 rpm by Celltron shaker (Infors HT, Switzerland) and were passaged every two to three days to 0.3-0.5 × 10^6^ cells/mL to maintain cells in the exponential phase. NucleoCounter®-250 (Chemometec, Denmark) was employed to determine cell density, viability and estimated cell diameter following the manufacturer’s instructions.

### Plasmid construction

Anti-apoptotic genes bcl-2 from human origin, bcl-xL from human origin, mcl-1 and bhrf-1 were codon optimised for *Cricetulus griseus* and supplied by GeneART™ (Thermo Fisher Scientific). Bcl-2 and bcl-xL from CHO origin (bcl-2 CHO and bcl-xL CHO) were extracted from cDNA from a CHO-S pellet harvested at mid-exponential phase. These DNA fragments were then inserted into a backbone plasmid via uracil-specific excision reagent (USER) cloning method (Lund et al., 2014).

The designed backbone plasmid had linker sequences for the USER cloning strategy followed by a polyA trap from bovine growth hormone (BGHpA) and separated from the EPO gene under RPU (relative promoter units) 100 promoter by 2 chicken b-globin locus control region hypersensitive sites 4 (HS4) insulators.

This construct was flanked by loxP (ATAACTTCGTATAGCATACATTATACGAAGTTAT) and lox2272 (ATAACTTCGTATAGGATACTTTATACGAAGTTAT). To facilitate colony selection and amplification in *E. coli* Mach1, the plasmid had the ampicillin resistance gene.

DNA bricks for USER assembly were generated by PCR amplification with Phusion U Hot Start DNA polymerase (Thermo Fisher Scientific) and uracil-containing primers (Integrated DNA Technologies, Coralville, IA, USA). Then, the product of the USER assembly was transformed into *E. coli* Mach1 competent cells (Thermo Fisher Scientific) for plasmid amplification. Sagner sequencing and restriction digestion was used for the verification of all constructs. All plasmid designs are available in Supplementary Figure S1.

### Cell line generation and verification

We developed all cell lines with relevant anti-apoptotic genes following the previously published process for cell line generation by our group (Grav et al., 2018). Briefly, the master cell line (MCL) had the mCherry gene flanked by loxP/lox2272 sites to allow for recombinase-mediated cassette exchange (RMCE) (Grav et al., 2018). The MCL was co-transfected with a plasmid containing EPO flanked by recombination sites loxP and lox2272 and a second plasmid with Cre-recombinase. Successfully recombinant clones switched the mCherry gene for EPO and were single-cell sorted into flat-bottom Corning 384-well plate (Sigma-Aldrich) containing 30 µL of CD CHO medium with 8 mM glutamine and 1.5% HEPES buffer (Gibco™) using the SH800S Cell Sorter (Sony biotechnology, CA, USA) based on mCherry^-^ phenotype. A negative control cell line was designed replacing the mCherry gene for 3 stop codons to ensure that the observed outcomes were not a product of the sorting process.

After single-cell sorting, the presence of only one cell per well was assessed with a Celigo Imaging Cell Cytometer (Nexcelom Bioscience). Ten to 14 days after the sorting, subconfluent clones were transferred to flat-bottom 96-well plates with 180 µL of CD CHO medium with 0.01% of Antibiotic-Antimycotic 100X and 8 mM of glutamine using an epMotion 5070 liquid handling workstation (Eppendorf). Subsequently, cells were expanded to flat-bottom 12-well plates and to 125-mL shake flasks for cell banking.

To ensure that gene expression only occurs with successful recombination, the promoter EF1α was located before the loxP site. A junction PCR was designed with primers binding at each end of the loxP/lox2272. Clones with a band corresponding to the expected DNA size were selected for the quantitative PCR (qPCR). In this second step, TaqMan Gene Expression Master Mix (Thermo Fisher Scientific), custom-made TaqMan assays for EPO and one-copy endogenous reference gene C1GALT1C1 (Cosmc) (Yang et al., 2014) were used in the duplex assay (EPO and Cosmc).

### BaxBak KO generation

All anti-apoptotic generated cell lines expressed bcl-2 family of proteins that prevent the pore forming activity of Bax and Bak on the mitochondrial membrane. Consequently, we knocked out (KO) the Bax and Bak genes from the negative control cell line, which contained 3 stop codons instead an anti-apoptotic gene to ensure that any observed differences were solely the consequence of the successful knock-out. For the generation of the BaxBak KO, the method based on CRISPR/Cas9 editing published by Grav et al. was followed (Grav et al., 2017). Briefly, specific single guide RNA (sgRNA) were designed for each gene (GGAAGCCGGTCAAACCACGTTGG for Bak and GCTGATGGCAACTTCAACTGGGG for Bax) using the CRISPy bioinformatics tool (Ronda et al., 2014).

The sgRNAs were synthesized, annealed, and cloned to an expression vector backbone harbouring scaffold RNA sequence, the U6 promoter and a termination sequence to generate sgRNA expression (Grav et al., 2017). The GFP 2A peptide-linked Cas9 (GFP_2A_Cas9) expression plasmid was used (Grav et al., 2015). A plasmid containing the sgRNA sequence of interest was co-transfected with the GFP_2A_Cas9 plasmid. Single cell sorting was performed to separate GFP positive cells. The KO was verified by Sagner sequencing and Western Blotting (Supplementary Figure S3).

### Sodium butyrate (NaBu) assay

In order to trigger apoptosis in the genetically engineer cell lines, cells were resuspended in CD CHO supplemented with 8 mM Glutamine, 1 % of Anti-Clumping Agent, 0.01 % of Antibiotic-Antimycotic and 20 mM sodium butyrate (Sigma-Aldrich - Merck Life Science B5887-1G LOT:SBLBQ8041V) to reach a final concentration of 2.5 × 10^6^ cells/mL. Then, cells were seeded in a 6-well plate with a working volume of 3 mL.

### Batch cultivation in 125-mL shake flask

Successfully generated cell lines were seeded at 0.5 × 10^6^ cells/mL in 125-mL shake flasks with 30 mL of CD CHO supplemented with 8 mM Glutamine, 1 % of Anti-Clumping Agent and 0.01 % of Antibiotic-Antimycotic for the batch characterisation.

Daily, 1 mL of cell culture was removed from each shake flask. 50 µL were utilized to measure cell density (cell/mL), viability (%) and estimated cell size (µm) with the NucleoCounter®-250. The remaining volume was centrifuged at 1000 g for 1 min and 400 µL of supernatant was run in the BioProfile FLEX2 (Nova Biomedical, Waltham, MA, USA) for glucose, glutamine, ammonia and glutamate measurements. The remaining volume from day 3 onwards was stored at -80 °C for subsequent EPO titer analysis with an Octet RED96 system (ForteBio, Sartorius, Göttingen, Germany). Finally, the pellet for day 3 and day 6 was stored at -80 °C for proteomics study.

Statistical analysis of EPO concentrations obtained from each cell line at day 6, 7 and 8 were performed with GraphPad Prism software (Version 10.02.01, GraphPad Software, San Diego, CA) using a Dunnett’s test with the negative control cell line as the control.

### Fed-batch in the ambr®15 system

Three cell lines were further characterised in a fed-batch culture was run in the ambr®15 system (Sartorius, Göttingen, Germany). Three bioreactors were inoculated for each cell line (CHO-S bcl-2 CHO, CHO-S bhrf-1 and CHO-S Negative Control) at 0.5 × 10^6^ cells/mL with 13 mL of 8 mM Glutamine, 1 % of Anti-Clumping Agent and 0.01 % of Antibiotic-Antimycotic. Daily, 0.2 µL of Anti-Clumping Reagent was added to each vessel to prevent foam accumulation.

### Proteomics study and statistical analysis

Protein identification was performed as described previously (Lavado-García et al., 2020). Briefly, MS/MS scans were matched against a *Cricetulus griseus* (UniProtKB/Swiss-Prot 2023_10 Release). Sequence of eGFP and Bhrf-1 were added to the previous database to enable their identification. For comparative analysis of changes in protein abundance, we applied weighted scan−peptide−protein statistical workflow, using SanXoT package (Martínez-Acedo et al., 2012; Navarro et al., 2014; Trevisan-Herraz et al., 2019).

The quantitative information is, as detailed by Lavado-García et al. 2020, obtained from the spectra and used to quantify the peptides from which the spectra are produced and then the proteins that generate these peptides (García-Marqués et al., 2016). These standardized variables (Zq) express the quantitative values in units of standard deviation (Navarro et al., 2014). The quantified proteins were functionally annotated using the Gene Ontology (GO) database. For further Gene Ontology annotation, the database for annotation, visualization and integrated discovery (DAVID) was used to perform functional enrichment analysis and to extract adjusted Benjamini scores for the enriched processes (Benjamini & Hochberg, 1995).

Specific consumption and production rates were calculated as indicated in Equation 1 where x_f_ equals final concentration of metabolite of interest, x_0_ equals initial concentration of metabolite of interest, t_f_ final time, t_0_ initial time, VCD_0_ viable cell density at initial time and VCD_f_ viable cell density at final time.

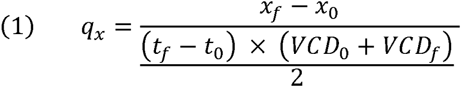

Considering that the sampling points for the proteomics analysis were day 3 and day 6, initial time equals to 3 and final time equals to 6 in the previous formula. Values were compared with a one-way ANOVA via GraphPad Prism software (Version 10.02.01, GraphPad Software, San Diego, CA). When the results of the analysis showed that the means were statistically significant, a Dunnet’s test was used to compare all means against the negative control.

## DATA AVAILABILITY

The raw mass spectrometry data has been submitted to the ProteomeXchange Consortium (http://proteomecentral.proteomexchange.org) via the PRIDE database with the dataset identifier PXD052672.

## AUTHOR CONTRIBUTIONS

DCT: Experimentation, investigation, visualization, writing original draft, review and editing. SG: experimentation. IMM and LG: experimental design and conceptualisation. JLG: Conceptualisation, visualization, supervision, review and editing. LKN: Supervision and review.

## ACKNOWLEDGMENTS

The authors would like to acknowledge generous support by Novo Nordisk Foundation. This work was supported by NNF20CC0035580 and NNF20SA0066621. LKN is supported by NNF14OC0009473. JLG is supported by NNF22OC0078741 and Marie Skłodowska-Curie Actions (MSCA) Postdoctoral Fellowship 101105465.

## REFERENCES

Altamirano, C., Paredes, C., Cairó, J. J., & Gòdia, F. (2000). Improvement of CHO cell culture medium formulation: Simultaneous substitution of glucose and glutamine. Biotechnology Progress, 16(1), 69–75. 10.1021/bp990124j

Benjamini, Y., & Hochberg, Y. (1995). Controlling the false discovery rate : A practical and powerful approach to multiple testing author (s): Yoav Benjamini and Yosef Hochberg Source : Journal of the Royal Statistical Society. Series B (Methodological), Vol. 57, No. 1 (1995), Publi. Journal of the Royal Statistical Society, 57(1), 289–300.

Bibila, T. A., & Robinson, D. K. (1995). In Pursuit of the Optimal Fed-Batch Process for Monoclonal Antibody Production. Biotechnology Progress, 11(1), 1–13. 10.1021/bp00031a001

Bielser, J. M., Wolf, M., Souquet, J., Broly, H., & Morbidelli, M. (2018). Perfusion mammalian cell culture for recombinant protein manufacturing – A critical review. Biotechnology Advances, 36(4), 1328–1340. 10.1016/j.biotechadv.2018.04.011

Clincke, M. F., Mölleryd, C., Zhang, Y., Lindskog, E., Walsh, K., & Chotteau, V. (2013). Very high density of CHO cells in perfusion by ATF or TFF in WAVE bioreactor^TM^: Part I: Effect of the cell density on the process. Biotechnology Progress, 29(3), 754–767. 10.1002/btpr.1704

Coulet, M., Kepp, O., Kroemer, G., & Basmaciogullari, S. (2022). Metabolic Profiling of CHO Cells during the Production of Biotherapeutics. Cells, 11(12), 1–21. 10.3390/cells11121929

Fussenegger, M., Fassnacht, D., Schwartz, R., Zanghi, J. A., Graf, M., Bailey, J. E., & Pörtner, R. (2000). Regulated overexpression of the survival factor bcl-2 in CHO cells increases viable cell density in batch culture and decreases DNA release in extended fixed-bed cultivation. Cytotechnology, 32(1), 45–61. 10.1023/A:1008168522385

García-Marqués, F., Trevisan-Herraz, M., Martínez-Martínez, S., Camafeita, E., Jorge, I., Lopez, J. A., Méndez-Barbero, N., Méndez-Ferrer, S., Del Pozo, M. A., Ibáñez, B., Andrés, V., Sánchez-Madrid, F., Redondo, J. M., Bonzon-Kulichenko, E., & Vázquez, J. (2016). A novel systems-biology algorithm for the analysis of coordinated protein responses using quantitative proteomics. Molecular and Cellular Proteomics, 15(5), 1740–1760. 10.1074/mcp.M115.055905

Grav, L. M., la Cour Karottki, K. J., Lee, J. S., & Kildegaard, H. F. (2017). Application of CRISPR/Cas9 genome editing to improve recombinant protein production in CHO cells. In Methods in Molecular Biology. 10.1007/978-1-4939-6972-2_7

Grav, L. M., Lee, J. S., Gerling, S., Kallehauge, T. B., Hansen, A. H., Kol, S., Lee, G. M., Pedersen, L. E., & Kildegaard, H. F. (2015). One-step generation of triple knockout CHO cell lines using CRISPR/Cas9 and fluorescent enrichment. Biotechnology Journal, 10(9), 1446–1456. 10.1002/biot.201500027

Grav, L. M., Sergeeva, D., Lee, J. S., Marin de Mas, I., Lewis, N. E., Andersen, M. R., Nielsen, L. K., Lee, G. M., & Kildegaard, H. F. (2018). Minimizing Clonal Variation during Mammalian Cell Line Engineering for Improved Systems Biology Data Generation. ACS Synthetic Biology, 7(9), 2148–2159. 10.1021/acssynbio.8b00140

Han, Y. K., Ha, T. K., Kim, Y.-G., & Lee, G. M. (2011). Bcl-xL overexpression delays the onset of autophagy and apoptosis in hyperosmotic recombinant Chinese hamster ovary cell cultures. Journal of Biotechnology, 156(1), 52–55. 10.1016/j.jbiotec.2011.07.032

Henry, M. N., MacDonald, M. A., Orellana, C. A., Gray, P. P., Gillard, M., Baker, K., Nielsen, L. K., Marcellin, E., Mahler, S., & Martínez, V. S. (2020). Attenuating apoptosis in Chinese hamster ovary cells for improved biopharmaceutical production. Biotechnology and Bioengineering, 117(4), 1187–1203. 10.1002/bit.27269

Joon, C. Y., Gatti, M. D. L., Philp, R. J., Yap, M., & Hu, W. S. (2008). Genomic and proteomic exploration of CHO and hybridoma cells under sodium butyrate treatment. Biotechnology and Bioengineering, 99(5), 1186–1204. 10.1002/bit.21665

Ko, P., Misaghi, S., Hu, Z., Zhan, D., Tsukuda, J., Yim, M., Sanford, M., Shaw, D., Shiratori, M., Snedecor, B., Laird, M., & Shen, A. (2018). Probing the importance of clonality: Single cell subcloning of clonally derived CHO cell lines yields widely diverse clones differing in growth, productivity, and product quality. Biotechnology Progress, 34(3), 624–634. 10.1002/btpr.2594

Krampe, B., & Al-Rubeai, M. (2010). Cell death in mammalian cell culture: Molecular mechanisms and cell line engineering strategies. Cytotechnology, 62(3), 175–188. 10.1007/s10616-010-9274-0

Kumar, S., Dhara, V. G., Orzolek, L. D., Hao, H., More, A. J., Lau, E. C., & Betenbaugh, M. J. (2021). Elucidating the impact of cottonseed hydrolysates on CHO cell culture performance through transcriptomic analysis. Applied Microbiology and Biotechnology, 105(1), 271–285. 10.1007/s00253-020-10972-7

Lavado-García, J., Jorge, I., Cervera, L., Vázquez, J., & Gòdia, F. (2020). Multiplexed Quantitative Proteomic Analysis of HEK293 Provides Insights into Molecular Changes Associated with the Cell Density Effect, Transient Transfection, and Virus-Like Particle Production. Journal of Proteome Research, 19(3), 1085–1099. 10.1021/acs.jproteome.9b00601

Lee, J. S., Kallehauge, T. B., Pedersen, L. E., & Kildegaard, H. F. (2015). Site-specific integration in CHO cells mediated by CRISPR/Cas9 and homology-directed DNA repair pathway. Scientific Reports, 5. 10.1038/srep08572

Lund, A. M., Kildegaard, H. F., Petersen, M. B. K., Rank, J., Hansen, B. G., Andersen, M. R., & Mortensen, U. H. (2014). A versatile system for USER cloning-based assembly of expression vectors for mammalian cell engineering. PLoS ONE, 9(5). 10.1371/journal.pone.0096693

MacDonald, M. A., Barry, C., Groves, T., Martínez, V. S., Gray, P. P., Baker, K., Shave, E., Mahler, S., Munro, T., Marcellin, E., & Nielsen, L. K. (2022). Modeling apoptosis resistance in CHO cells with CRISPR-mediated knockouts of Bak1, Bax, and Bok. Biotechnology and Bioengineering, 119(6), 1380–1391. 10.1002/bit.28062

Majors, B. S., Betenbaugh, M. J., Pederson, N. E., & Chiang, G. G. (2008). Enhancement of transient gene expression and culture viability using Chinese hamster ovary cells overexpressing Bcl-xL. Biotechnology and Bioengineering, 101(3), 567–578. 10.1002/bit.21917

Majors, B. S., Betenbaugh, M. J., Pederson, N. E., & Chiang, G. G. (2009). Mcl-1 overexpression leads to higher viabilities and increased production of humanized monoclonal antibody in Chinese hamster ovary cells. Biotechnology Progress, 25(4), 1161–1168. 10.1002/btpr.192

Martínez-Acedo, P., Núñez, E., Sánchez Gómez, F. J., Moreno, M., Ramos, E., Izquierdo-Álvarez, A., Miró-Casas, E., Mesa, R., Rodriguez, P., Martínez-Ruiz, A., Dorado, D. G., Lamas, S., & Vázquez, J. (2012). A novel strategy for global analysis of the dynamic thiol redox proteome. Molecular and Cellular Proteomics, 11(9), 800–813. 10.1074/mcp.M111.016469

Martínez-Monge, I., Comas, P., Triquell, J., Casablancas, A., Lecina, M., Paredes, C. J., & Cairó, J. J. (2019). Concomitant consumption of glucose and lactate: A novel batch production process for CHO cells. Biochemical Engineering Journal, 151(April), 107358. 10.1016/j.bej.2019.107358

Martínez, V. S., Dietmair, S., Quek, L., Hodson, M. P., Gray, P., & Nielsen, L. K. (2013). Flux balance analysis of CHO cells before and after a metabolic switch from lactate production to consumption. Biotechnology and Bioengineering, 110(2), 660–666. 10.1002/bit.24728

Mastrangelo, A. J., Hardwick, J. M., Bex, F., & Betenbaugh, M. J. (2000). Part I. Bcl-2 and Bcl-X(L) limit apoptosis upon infection with alphavirus vectors. Biotechnology and Bioengineering, 67(5), 544–554. 10.1002/(sici)1097-0290(20000305)67:5<544::aid-bit5>3.0.co;2-%23

Mohamed, A.-R., Rabinder, P. S., Al-Rubeai, M., Singh, R. P., Mohamed, A.-R., & Rabinder, P. S. (1998). Apoptosis in cell culture. Current Opinion in Biotechnology, 9(2), 152–156. 10.1016/S0958-1669(98)80108-0

Navarro, P., Trevisan-Herraz, M., Bonzon-Kulichenko, E., Núñez, E., Martínez-Acedo, P., Pérez-Hernández, D., Jorge, I., Mesa, R., Calvo, E., Carrascal, M., Hernáez, M. L., García, F., Bárcena, J. A., Ashman, K., Abian, J., Gil, C., Redondo, J. M., & Vázquez, J. (2014). General statistical framework for quantitative proteomics by stable isotope labeling. Journal of Proteome Research, 13(3), 1234–1247. 10.1021/pr4006958

O’Flaherty, R., Bergin, A., Flampouri, E., Mota, L. M., Obaidi, I., Quigley, A., Xie, Y., & Butler, M. (2020). Mammalian cell culture for production of recombinant proteins: A review of the critical steps in their biomanufacturing. Biotechnology Advances, 43(April), 107552. 10.1016/j.biotechadv.2020.107552

Ochoa, S. (2016). A new approach for finding smooth optimal feeding profiles in fed-batch fermentations. Biochemical Engineering Journal, 105, 177–188. 10.1016/j.bej.2015.09.004

Rahimi, A., Karimipoor, M., Mahdian, R., Alipour, A., Hosseini, S., Mohammadi, M., Kaghazian, H., Abbasi, A., Shahsavarani, H., & Shokrgozar, M. A. (2023). Efficient CRISPR/Cas9-Mediated BAX Gene Ablation in CHO Cells To Impair Apoptosis and Enhance Recombinant Protein Production. Iranian Journal of Biotechnology, 21(2), 75–86. 10.30498/ijb.2023.343428.3388

Ronda, C., Pedersen, L. E., Hansen, H. G., Kallehauge, T. B., Betenbaugh, M. J., Nielsen, A. T., & Kildegaard, H. F. (2014). Accelerating genome editing in CHO cells using CRISPR Cas9 and CRISPy, a web-based target finding tool. Biotechnology and Bioengineering, 111(8), 1604–1616. 10.1002/bit.25233

Singh, R., Letai, A., & Sarosiek, K. (2019). Regulation of apoptosis in health and disease: the balancing act of BCL-2 family proteins. Nature Reviews Molecular Cell Biology, 20(3), 175–193. 10.1038/s41580-018-0089-8

Singh, R. P., Al.-Rubeai, M., Gregory, C. D., & Emery, A. N. (1994). Cell death in bioreactors: A role for apoptosis. Biotechnology and Bioengineering, 44(6), 720–726. 10.1002/bit.260440608

Templeton, N., Lewis, A., Dorai, H., Qian, E. A., Campbell, M. P., Smith, K. D., Lang, S. E., Betenbaugh, M. J., & Young, J. D. (2014). The impact of anti-apoptotic gene Bcl-2Δ expression on CHO central metabolism. Metabolic Engineering, 25, 92–102. 10.1016/j.ymben.2014.06.010

Trevisan-Herraz, M., Bagwan, N., García-Marqués, F., Rodriguez, J. M., Jorge, I., Ezkurdia, I., Bonzon-Kulichenko, E., & Vázquez, J. (2019). SanXoT: a modular and versatile package for the quantitative analysis of high-throughput proteomics experiments. Bioinformatics, 35(9), 1594–1596. 10.1093/bioinformatics/bty815

Wahl, M. C., Will, C. L., & Lührmann, R. (2009). The Spliceosome: Design Principles of a Dynamic RNP Machine. Cell, 136(4), 701–718. 10.1016/j.cell.2009.02.009

Wang, M. D., Yang, M., Huzel, N., & Butler, M. (2002). Erythropoietin production from CHO cells grown by continuous culture in a fluidized-bed bioreactor. Biotechnology and Bioengineering, 77(2), 194–203. 10.1002/bit.10144

Wong, D. C. F., Wong, K. T. K., Lee, Y. Y., Morin, P. N., Heng, C. K., & Yap, M. G. S. (2006). Transcriptional profiling of apoptotic pathways in batch and fed-batch CHO cell cultures. Biotechnology and Bioengineering, 94(2), 373–382. 10.1002/bit.20872

Xiong, K., Marquart, K. F., la Cour Karottki, K. J., Li, S., Shamie, I., Lee, J. S., Gerling, S., Yeo, N. C., Chavez, A., Lee, G. M., Lewis, N. E., & Kildegaard, H. F. (2019). Reduced apoptosis in Chinese hamster ovary cells via optimized CRISPR interference. Biotechnology and Bioengineering, 116(7), 1813–1819. 10.1002/bit.26969

Yang, Z., Halim, A., Narimatsu, Y., Jitendra Joshi, H., Steentoft, C., Gram Schjoldager, K. T.-B., Alder Schulz, M., Sealover, N. R., Kayser, K. J., Paul Bennett, E., Levery, S. B., Vakhrushev, S. Y., & Clausen, H. (2014). The GalNAc-type O-Glycoproteome of CHO Cells Characterized by the SimpleCell Strategy. Molecular & Cellular Proteomics, 13(12), 3224–3235. 10.1074/mcp.M114.041541

Yu, S., Meng, S., Xiang, M., & Ma, H. (2021). Phosphoenolpyruvate carboxykinase in cell metabolism: Roles and mechanisms beyond gluconeogenesis. Molecular Metabolism, 53(May), 101257. 10.1016/j.molmet.2021.101257

