## Supplementary Figures for "Evaluating apoptotic gene efficiency for CHO culture performance using targeted integration"

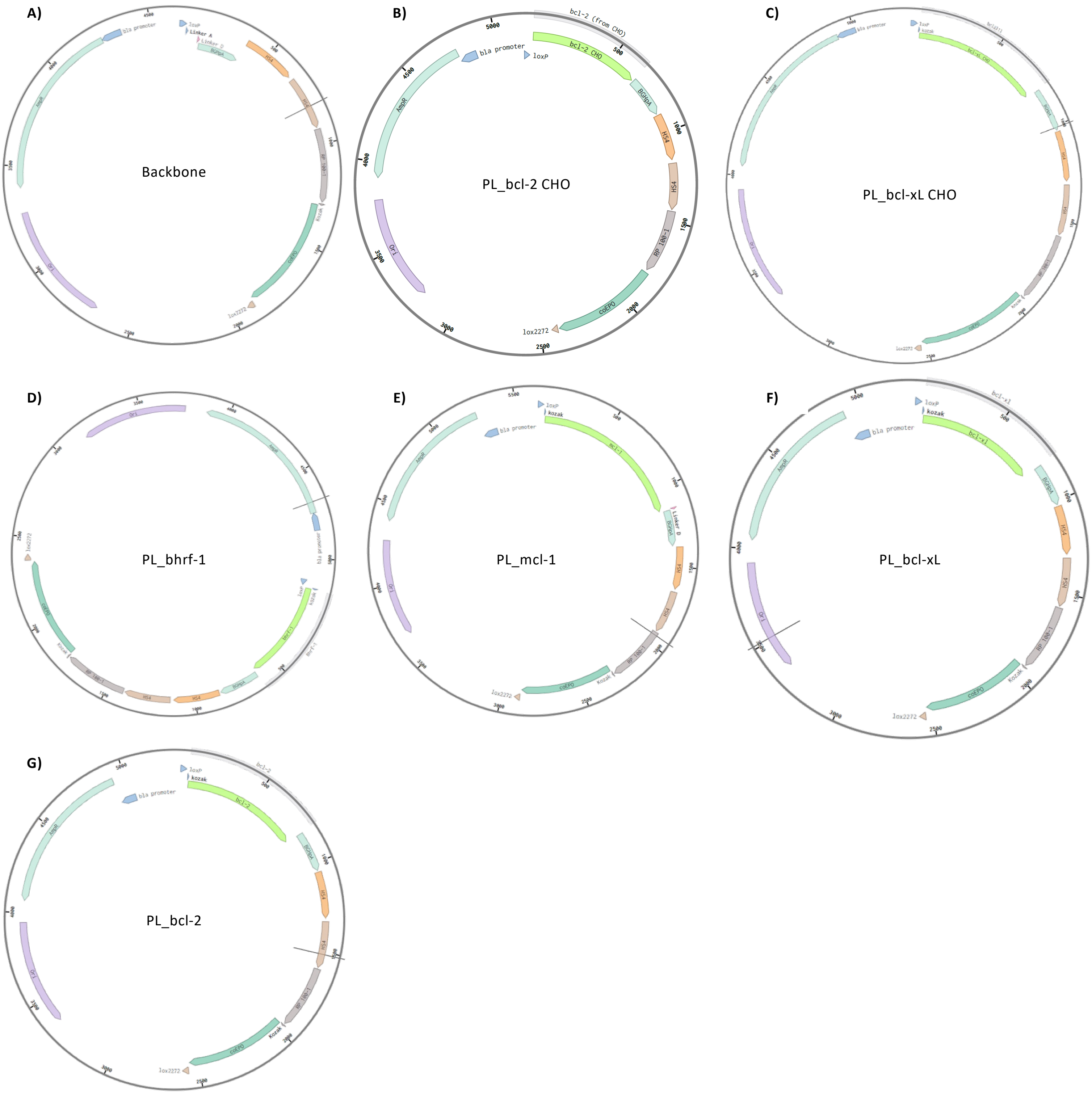


**Supplementary Figure S1:** (A) Backbone used to introduce all anti-apoptotic genes studied using the linker strategy. (B), (C), (D), (E), (F), and (G) illustrate the final plasmid originated from the backbone and transfected to generate the isogenic-stable cell lines.

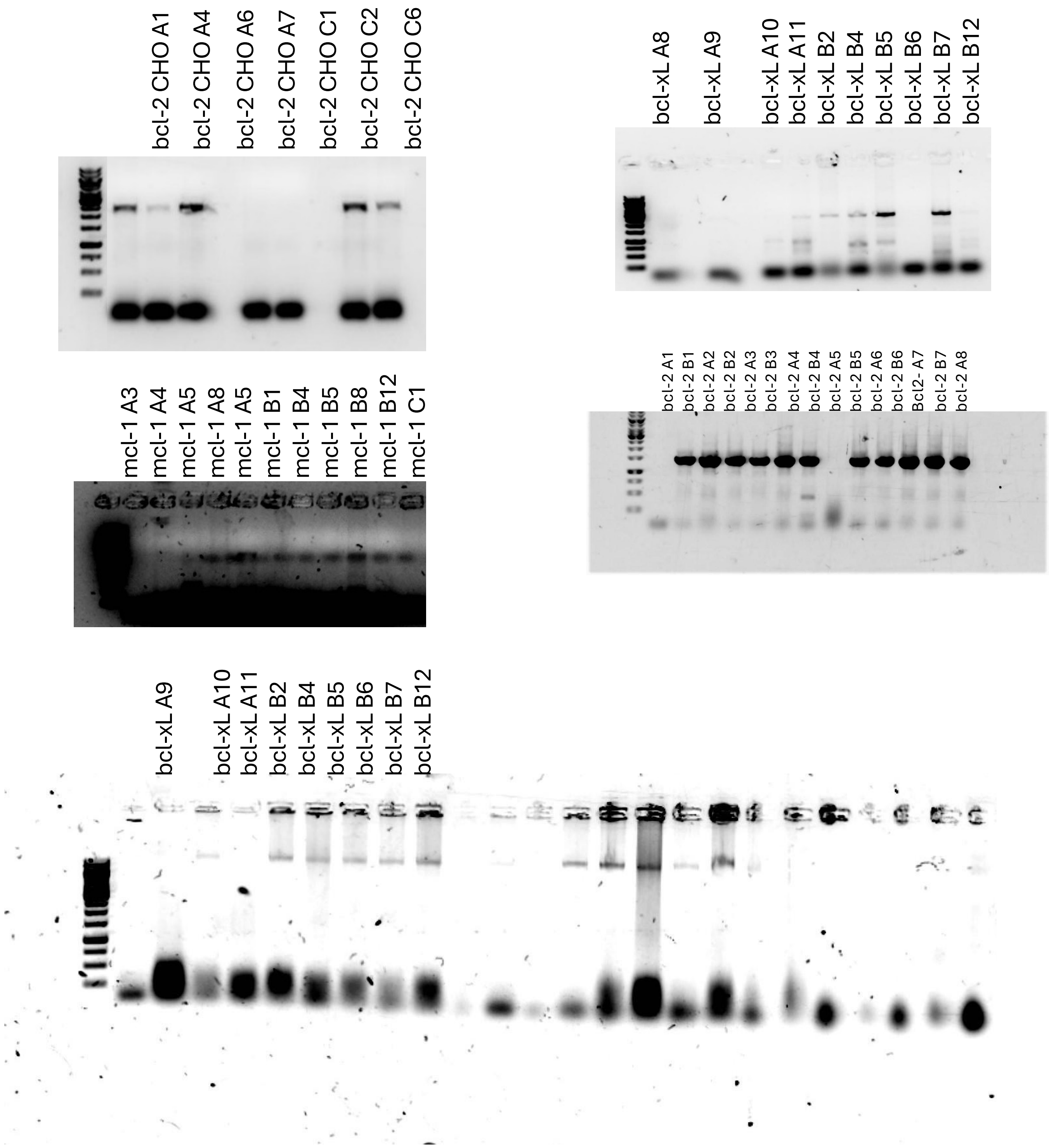


**Supplementary Figure S2:** Junction PCR results. The selected clones for all experiments are marked in red.

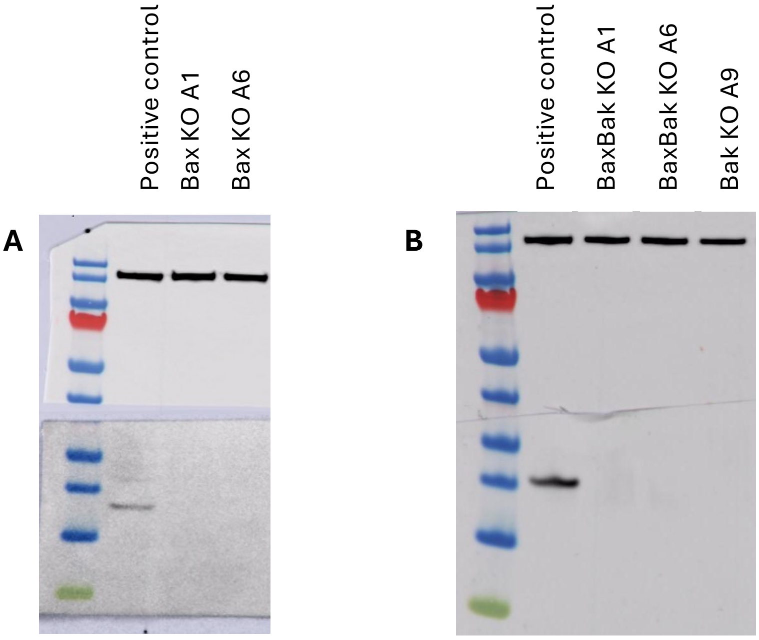


**Supplementary Figure S3:** Western Blotting to verify the successful knockout of (A) Bax (23 kDa) and then (B) Bak (20 kDa) with Vinculin (143 kDa) as housekeeping gene.

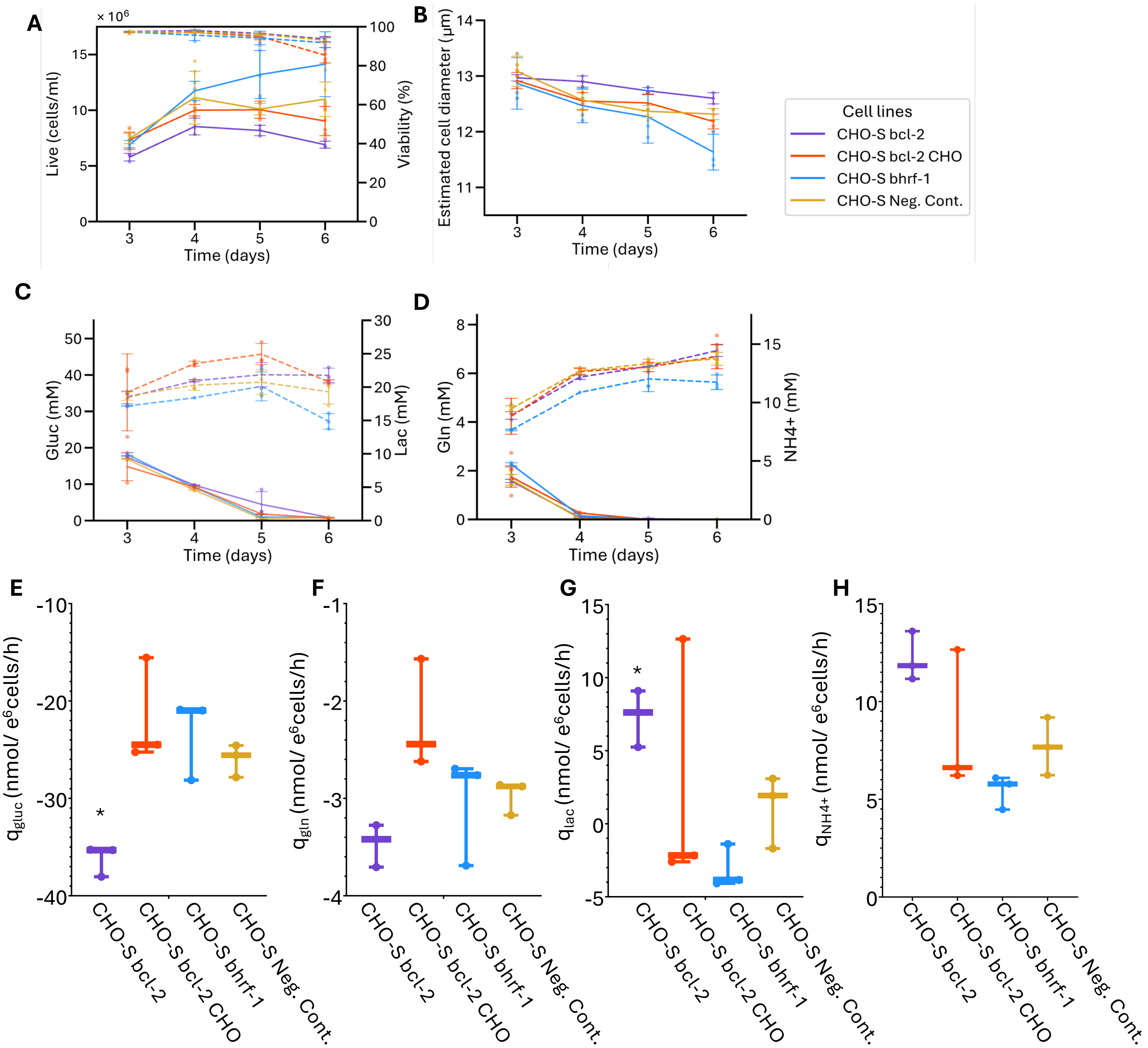


**Supplementary Figure 4:** Development from day 3 to day 6 of: (A) viable cell density in cells/mL, (B) viability in %, (C) glucose and lactate concentration in mM and (D) glutamine and ammonia concentration in mM. (E) glucose consumption rate (nmol/e[6]cells/h), (F) glutamine consumption rate (nmol/e[6]cells/h), (G) lactate consumption/production rate (nmol/e[6]cells/h), (H) ammonia production rate (nmol/e[6]cells/h). Statistically significant values compared to the negative control with a Dunnett's test are (p-value<0.05) illustrated with "*".
